# Network analysis of genome-wide selective constraint reveals a gene network active in early fetal brain intolerant of mutation

**DOI:** 10.1101/017277

**Authors:** Jinmyung Choi, Parisa Shooshtari, Kaitlin E Samocha, Mark J Daly, Chris Cotsapas

## Abstract

Using robust, integrated analysis of multiple genomic datasets, we show that genes depleted for non-synonymous *de novo* mutations form a subnetwork of 72 members under strong selective constraint. We further show this subnetwork is preferentially expressed in the early development of the human hippocampus and is enriched for genes mutated in neurological, but not other, Mendelian disorders. We thus conclude that carefully orchestrated developmental processes are under strong constraint in early brain development, and perturbations caused by mutation have adverse outcomes subject to strong purifying selection. Our findings demonstrate that selective forces can act on groups of genes involved in the same process, supporting the notion that adaptation can act coordinately on multiple genes. Our approach provides a statistically robust, interpretable way to identify the tissues and developmental times where groups of disease genes are active. Our findings highlight the importance of considering the interactions between genes when analyzing genome-wide sequence data.

## Introduction

Genetic variation is introduced into the human genome by spontaneously arising *de novo* mutations in the germline. The majority of these mutations have, at most, modest effects on phenotype; they are thus subject to nearly neutral drift and can be transmitted through the population, with some increasing in frequency to become common variants. Conversely, *de novo* mutations with large effects on phenotype are subject to many different selective forces, both positive and negative, with the latter resulting in either the variant being completely lost from the population or maintained at very low frequencies^1^.

Large-scale DNA sequencing can now be used to comprehensively assess *de novo* mutations, with current applications focusing on the protein-coding portion of the genome (the exome). This approach has been used to identify causal genes and variants in rare mendelian diseases: for example, exome sequencing of ten affected individuals with Kabuki syndrome identified the methyl transferase KMT2D (formerly MLL2) as causal, after substantial *post hoc* data filtering^2^. In complex traits, this approach has successfully identified pathogenic genes harboring *de novo* mutations in autism spectrum disorders, intellectual disability and two epileptic encephalopthies^3^; notably, all these studies sequenced the exomes of parent-affected offspring trios and quantified the background rate of *de novo* mutations in each gene using formal analytical approaches. They were thus able to identify genes harboring a statistically significant number of mutations, which are likely to be causal for disease^3, 4^.

These large-scale exome sequencing studies have demonstrated that the rate of non-synonymous *de novo* mutations is markedly depleted in some genes, and that these genes harbor disease-causing mutations. As synonymous *de novo* mutations occur at expected frequencies, this depletion is not driven by variation in the local overall mutation rate; instead, these genes appear to be intolerant of changes to amino acid sequence and are thus under selective constraint, with non-synonymous mutations removed by purifying selection. These genes represent a limited number of fundamental biological roles, which suggests that entire processes, rather than single genes, are under selective constraint. This is consistent with the extreme polygenicity of most human traits, where hundreds of genes play a causal role in determining organismal phenotype^5, 6^. These genes must participate in the same cellular processes, but uncovering the relevant connections and the cell populations and developmental stages in which they occur remains a challenge. We and others have described statistical frameworks to test connectivity within a nominated set of genes^3, 4^; whilst these approaches are adequate for testing limited gene sets, there is still a dearth of systematic ways to assess connectivity in a genome-wide fashion and identify the tissues in which connected groups of genes are likely to act in a statistically rigorous and interpretable way.

We have developed a robust, unbiased framework to address these questions and applied it to genome-wide selective constraint data derived from exome sequences of 6,503 individuals^4^. We identified a single, statistically significant subnetwork of 72 interacting genes highly intolerant of non-synonymous variation, with no other interacting groups of genes showing evidence of such coordinate constraint. To establish biological context for this subnetwork, we developed a robust approach to test for preferential expression of the module as a whole, rather than the individual constituent genes. Using gene expression data from the cosmopolitan atlas of tissues in the Roadmap Epigenome Project^7, 8^, we found that this subnetwork is preferentially expressed in several early-stage tissues, with the strongest enrichment in fetal brain. To more carefully dissect the role of this subnetwork in the central nervous system, we analyzed expression data from BrainSpan^9^, an atlas of the developing human brain, and found that the constrained gene subnetwork is preferentially expressed in the early development of the hippocampus. Consistent with this observation, this module is enriched for genes mutated in neurological, but not other, Mendelian disorders. We thus show that selective constraint acts on a set of interacting genes active in early brain development, and that these genes are in fact intolerant of mutation. Our **P**rotein **I**nteraction **N**etwork **T**issue **S**earch (PINTS) framework is publicly available at https://www.dropbox.com/sh/hgwmf1qx3a5wdxz/AACUvEH4EAb3yKLnxKBAg_nxa?dl=0.

## Results

### Calculating selective constraint scores

We have previously described a framework to assess selective constraint across coding sequences in the genome^4^. Briefly, we calibrated an expectation for all possible conversions of one base to another by mutation from non-coding sequence. For each transition, we modeled the effect of the surrounding sequence and its conservation across species to correct for context effects. We then counted the number of synonymous and non-synonymous variants in the coding sequence of each gene in the genome and derived a statistic of constraint on each class of variation compared to this global expectation. We found that a number of genes show decreased rates of non-synonymous substitution but expected rates of synonymous substitution, consistent with purifying selection removing the non-synonymous alleles from the population.

### Analysis framework description

If constrained genes lie in biologically meaningful networks, we expect them to (i) interact and (ii) be expressed in the same tissues. We developed a robust, modular workflow (**PINTS** – **P**rotein **I**nteraction **N**etwork **T**issue **S**earch; Figure 1) to test both of these hypotheses at a genome-wide level. To detect interactions between constrained genes we used a high-confidence protein-protein interaction network (InWeb^10^), and employed a clustering algorithm previously validated on such networks^11^. We assessed significance empirically by randomly reassigning constraint scores to genes (see Methods and Supplementary material). We then tested any significant subnetworks for preferential expression in the diverse tissue atlas provided by the Roadmap Epigenome Project (REP), which assays gene expression in 27 human primary samples across the developmental spectrum^8^. Our final dataset is comprised of 9729 genes both present in InWeb and detected in at least one REP tissue.

**Figure 1:**
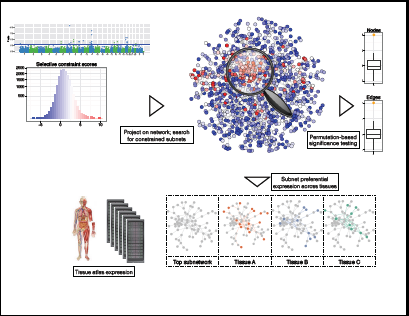
the Protein Interaction Network Tissue Search (PINTS) workflow. We project gene-wise selective constraint scores^4^ onto the InWeb protein-protein interaction dataset^10^ and use a heuristic version of the prize-collecting Steiner Tree algorithm^11, 18^ to detect clusters of interacting constrained genes. We assess significance empirically, by randomly assigning the scores to genes 1000 times and calibrating detected subnetwork parameters. We then test any significant subnetwork for usual patterns of preferential expression^21^ across the Roadmap Epigenome Project expression data^8^, a cosmopolitan tissue atlas, using a Markov random field approach. The approach is flexible and modular, so gene interaction and tissue expression reference datasets can be altered according to the application.

Our workflow is both modular and flexible: clustering algorithms, gene-gene relationships and tissue atlases can be replaced as required, so that analyses can be tailored to suit specific biological problems. A flexible implementation, including all data described here, is freely available as an R package at https://www.dropbox.com/sh/hgwmf1qx3a5wdxz/AACUvEH4EAb3yKLnxKBAg_nxa?dl=0.

### Highly constrained genes form a protein interaction module expressed in fetal tissues and the immune system

We define highly constrained genes as those with evidence of constraint on non-synonymous *de novo* substitutions (*p < 5 × 10*^-6^, Bonferroni correction for the number of genes in our InWeb dataset) but null synonymous constraint scores, indicating intolerance to functionally relevant mutation rather than fluctuations in the local mutation rate^4^. Of these, 107/9729 genes pass this stringent threshold (*p < 2.2 × 10^-16^*; Table S1), and form the core of the analysis presented here. We found that 67/107 form a connected subnetwork (Figure 2A; Table 1). Five additional genes are included as our cluster detection algorithm by design looks for a backbone of null nodes connected to many signal nodes. To assess the significance of this observation, we randomly distribute constraint scores to InWeb nodes 1000 times and find that the constrained subnetwork is larger (*p < 0.001*) and more densely connected (number of edges: *p < 0.001*; clustering coefficient: *p = 0.008*) than expected by chance (Figure 2B). As such, it also explains more total constraint in the genome than expected (*p < 0.001*). After accounting for the genes forming this subnetwork, we found no evidence of other independent subnetworks of constrained genes.

**Table 1:** a 72-member constrained gene subnetwork. We find that 67/107 significantly constrained genes form a single protein-protein interaction subnetwork. Five additional genes are also included (gray shading), as our cluster detection algorithm by design looks for a backbone of null nodes connected to many signal nodes. As shown in Figure 2, the subnetwork is significantly larger and more densely connected than expected by chance, and is preferentially expressed in a subset of early-stage neural and immune tissues.

| Gene | Constraint score | Chr | Start | End | Gene | Constraint score | Chr | Start | End |
| --- | --- | --- | --- | --- | --- | --- | --- | --- | --- |
| DYNC1H1 | 9.977 | 14 | 101964528 | 102050792 | UBR4 | 4.940 | 1 | 19074506 | 19210276 |
| PRPF8 | 8.302 | 17 | 1650629 | 1684882 | CHD3 | 4.905 | 17 | 7884806 | 7912760 |
| HUWE1 | 7.973 | X | 53532096 | 53686729 | USP7 | 4.866 | 16 | 8892094 | 8964514 |
| SMARCA4 | 6.604 | 19 | 10961001 | 11065395 | PRPF6 | 4.826 | 20 | 63981135 | 64033100 |
| POLR2A | 6.578 | 17 | 7484366 | 7514618 | GNAS | 4.806 | 20 | 58839718 | 58911192 |
| RYR2 | 6.436 | 1 | 237042205 | 237833988 | THOC2 | 4.791 | X | 123600561 | 123733056 |
| MED12 | 6.388 | X | 71118556 | 71142454 | FRY | 4.772 | 13 | 32031300 | 32299122 |
| SNRNP200 | 6.166 | 2 | 96274336 | 96305515 | OGT | 4.753 | X | 71533083 | 71575897 |
| CHD4 | 6.162 | 12 | 6570083 | 6607476 | POLR2B | 4.729 | 4 | 56977722 | 57031168 |
| MTOR | 5.974 | 1 | 11106535 | 11262507 | KCNMA1 | 4.687 | 10 | 76869601 | 77638595 |
| GRIN1 | 5.971 | 9 | 137138390 | 137168762 | TAOK1 | 4.685 | 17 | 29390464 | 29551904 |
| PPFIA3 | 5.794 | 19 | 49119389 | 49151026 | BRWD3 | 4.683 | X | 80670854 | 80809688 |
| MLL | 5.747 | 11 | 118436490 | 118526832 | SPTAN1 | 4.671 | 9 | 128552558 | 128633665 |
| UBR5 | 5.720 | 8 | 102253012 | 102412841 | PHIP | 4.670 | 6 | 78935867 | 79078236 |
| ITPR1 | 5.589 | 3 | 4493348 | 4847840 | DDB1 | 4.670 | 11 | 61299451 | 61342596 |
| CLTC | 5.547 | 17 | 59619689 | 59696956 | HSPA2 | 4.665 | 14 | 64535905 | 64546173 |
| FLNA | 5.541 | X | 154348524 | 154374638 | SPEG | 4.644 | 2 | 219434846 | 219498287 |
| UPF1 | 5.514 | 19 | 18831938 | 18868236 | SMC3 | 4.639 | 10 | 110567691 | 110604636 |
| HCFC1 | 5.450 | X | 153947553 | 153971807 | MYH10 | 4.629 | 17 | 8474205 | 8630761 |
| DHX30 | 5.428 | 3 | 47802909 | 47850195 | XPO1 | 4.621 | 2 | 61477849 | 61538626 |
| SPTBN1 | 5.423 | 2 | 54456285 | 54671445 | CUL3 | 4.610 | 2 | 224470150 | 224585397 |
| SF3B1 | 5.418 | 2 | 197389784 | 197435091 | IRS2 | 4.592 | 13 | 109752698 | 109786568 |
| SMARCA2 | 5.387 | 9 | 2015219 | 2193624 | ADCY1 | 4.587 | 7 | 45574140 | 45723116 |
| CACNA1I | 5.363 | 22 | 39570753 | 39689737 | APC2 | 4.564 | 19 | 1446302 | 1473244 |
| SMC1A | 5.360 | X | 53374149 | 53422728 | ZBTB17 | 4.547 | 1 | 15941869 | 15976132 |
| GRIN2B | 5.334 | 12 | 13537337 | 13980119 | TLN1 | 4.517 | 9 | 35696948 | 35732395 |
| GRIN2D | 5.211 | 19 | 48394875 | 48444931 | MYH9 | 4.496 | 22 | 36281281 | 36388018 |
| TAF1 | 5.178 | X | 71366239 | 71532374 | EEF2 | 4.478 | 19 | 3976056 | 3985469 |
| VCP | 5.162 | 9 | 35056064 | 35073249 | PDS5A | 4.451 | 4 | 39822863 | 39977956 |
| CNOT1 | 5.146 | 16 | 58519951 | 58629886 | PRKD2 | 4.438 | 19 | 46674275 | 46717127 |
| TRIO | 5.109 | 5 | 14143702 | 14532128 | BRD4 | 4.436 | 19 | 15235519 | 15332545 |
| CYFIP2 | 5.100 | 5 | 157266079 | 157395598 | HSPA8 | 4.364 | 11 | 123057489 | 123063230 |
| SUPT5H | 5.065 | 19 | 39436156 | 39476670 | CTNNB1 | 4.198 | 3 | 41194837 | 41260096 |
| FZD8 | 5.028 | 10 | 35638249 | 35642278 | UBC | 3.997 | 12 | 124911604 | 124917368 |
| TNPO2 | 4.993 | 19 | 12699194 | 12724011 | PIK3CD | 3.858 | 1 | 9651732 | 9729114 |
| GTF2I | 4.945 | 7 | 74657667 | 74760692 | PIK3R1 | 2.170 | 5 | 68215720 | 68301821 |

**Figure 2:**
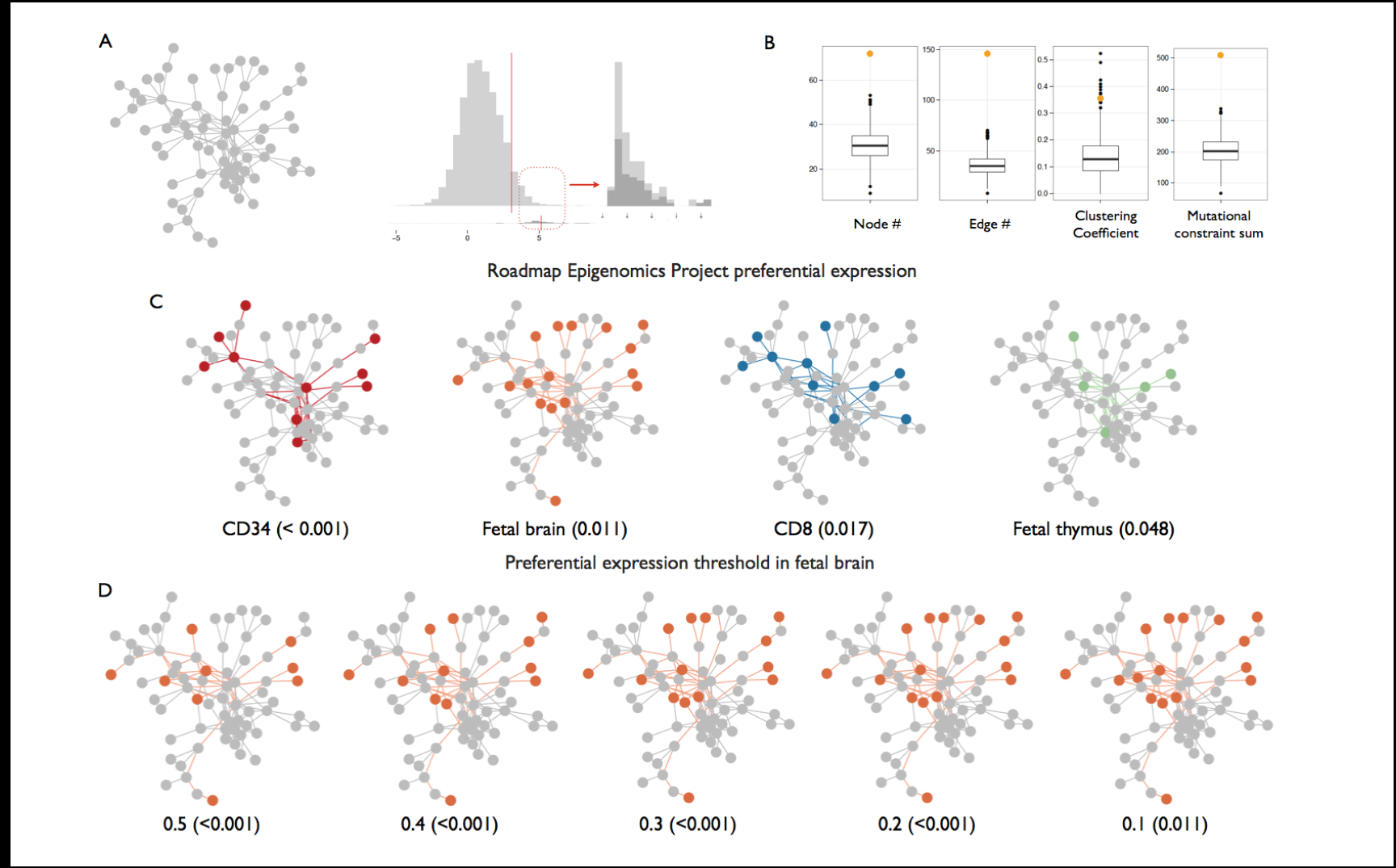
selectively constrained genes form a 72-member network, preferentially expressed in fetal brain, heart and immune cell populations. A: constrained genes form a connected subnetwork of genes in the extreme of the constraint score distribution. B: the constrained subnetwork contains more genes (node *p < 0.001*), has more connections (edge *p < 0.001*), is more densely connected (clustering coefficient *p = 0.008*) and explains more total constraint (sum *p < 0.001*) than expected by chance (orange dots) compared to networks discovered in 1000 permutations of the constraint data (boxplots and black dots). C: the constrained subnetwork is preferentially expressed in a subset of Roadmap Epigenome Project tissues, including fetal brain. D: The most consistent preferential expression signal is seen in fetal brain, which is robust to stringency of preferential expression threshold.

The genes in the constrained subnetwork appear to represent several fundamental cell processes, most notably mitosis and cell proliferation (SMC1A, SMC3, CTNNB1) and transcriptional regulation (CHD3, CHD4, SMARCA4). We performed a formal pathway analysis to further test this and found enrichment of several annotated pathways reflecting these fundamental processes (Table 2). Encouraged that our detected subnetwork represents one or more biological processes under constraint, we sought to add cellular context to our observations. In particular, we wanted to determine if this group of genes is preferentially expressed in particular tissues, indicating a likely site of action. We thus developed an approach to estimate the joint probability of preferential expression of the genes in the subnetwork in each tissue of an atlas of expression data, while accounting for how frequently each gene is detected across the entire atlas. We applied our approach, which uses Markov random fields, to the expression data on 27 primary tissues and cell lines available from the Roadmap Epigenome Project. Using two conservative permutation-based significance tests, we find the constrained subnetwork is preferentially expressed in a number of fetal and immune tissues (Figure 2C and Table 3), including fetal brain (*p < 0.001*), the immune cell subpopulations marked by CD34 (*p < 0.001*) and CD8 (p = 0.017) and fetal thymus (p = 0.048). We note that, whilst only a subset of genes are expressed in any one tissue, the combinations of genes expressed in these tissues is highly statistically significant: each gene is only expressed in a small subset of the tissues interrogated, so the cumulative probability of seeing these genes coordinately expressed in any one tissue is small.

**Table 2:** the 72-member constrained gene subnetwork is enriched for canonical pathways reflecting neuronal and immune functionality and basic aspects of cell cycle control. We tested pathways from two sources (the Reactome database and KEGG, the Kyoto Encyclopedia of Genes and Genomes), assessing how many genes are in each pathway (All), how many map onto the 9729 inteconnected genes in our analysis (Mapped), and how many are present in the constrained subnetwork (Subnetwork). We assess significance using both the GSEA approach of a Kolmogorov-Smirnov (KS) test and a simple hypergeometric (HG) test of expected overlaps.

| Name | All | Mapped | Subnetwork | KS | HG |
| --- | --- | --- | --- | --- | --- |
| Developmental biology (Reactome) | 397 | 344 | 10 | 2.53E-19 | 1.83E-05 |
| Immune system (Reactome) | 934 | 702 | 9 | 4.98E-08 | 1.96E-02 |
| Adaptive immune system (Reactome) | 540 | 421 | 8 | 3.13E-10 | 2.08E-03 |
| Axon guidance (Reactome) | 252 | 220 | 8 | 4.62E-16 | 1.65E-05 |
| mRNA Processing (Reactome) | 162 | 120 | 8 | 9.06E-12 | 1.05E-07 |
| Calcium signaling pathway (KEGG) | 179 | 163 | 7 | 6.00E-08 | 1.36E-05 |
| Spliceosome (KEGG) | 129 | 85 | 7 | 5.32E-21 | 9.60E-08 |
| mRNA splicing (Reactome) | 112 | 74 | 7 | 6.26E-20 | 3.19E-08 |
| Processing of capped intron containing pre-mRNA (Reactome) | 141 | 102 | 7 | 3.21E-13 | 3.99E-07 |
| Pathways in cancer (KEGG) | 329 | 301 | 6 | 4.05E-12 | 4.21E-03 |
| Regulation of actin cytoskeleton (KEGG) | 217 | 188 | 6 | 8.08E-13 | 2.74E-04 |
| Cell cycle (Reactome) | 422 | 332 | 6 | 7.95E-03 | 7.12E-03 |
| mRNA splicing minor pathway (Reactome) | 46 | 20 | 6 | 2.90E-05 | 3.56E-11 |
| Signalling by NGF (Reactome) | 218 | 191 | 6 | 3.04E-21 | 3.02E-04 |
| Focal adhesion (KEGG) | 202 | 188 | 5 | 9.72E-07 | 1.70E-03 |
| Long term potentiation (KEGG) | 71 | 60 | 5 | 9.64E-15 | 2.97E-06 |
| MAPK signaling pathway (KEGG) | 268 | 233 | 5 | 3.02E-15 | 4.92E-03 |
| HIV infection (Reactome) | 208 | 163 | 5 | 1.32E-06 | 8.12E-04 |
| HIV life cycle (Reactome) | 126 | 95 | 5 | 1.03E-02 | 4.27E-05 |
| Late phase of HIV life cycle (Reactome) | 105 | 85 | 5 | 1.36E-02 | 2.27E-05 |
| Neuronal system (Reactome) | 280 | 219 | 5 | 3.80E-28 | 3.64E-03 |
| NGF signalling via TRKa from the plasma membrane (Reactome) | 138 | 120 | 5 | 1.09E-14 | 1.57E-04 |
| Signaling by GPCR (Reactome) | 921 | 415 | 5 | 1.63E-11 | 6.21E-02 |

As several tissues are enriched for subnetwork expression, we sought to understand whether we were capturing the same signature across multiple tissues reflecting a shared process. We assessed whether the same genes are preferentially expressed in each tissue, and found a distinct signature in the fetal brain and heart samples and the immune cell subpopulations (CD34, CD8, CD3, thymus; pairwise *p < 0.05* hypergeometric test; Table S2). To ensure our tissue expression results are not an artifact of the threshold we set for preferential expression, we repeated the entire analysis with a range of threshold values and found consistent results across tissues; this is most notable in fetal brain (Figure 2D and Table S3), which remains significant irrespective of threshold used.

Genes under selective constraint are more likely to harbor pathogenic mutations causing mendelian diseases, consistent with intolerance of functional mutations^4^. Accordingly, we found that our subnetwork of 72 genes is significantly enriched for OMIM annotations (*p = 0.0013*). To further elucidate this observation, we mapped all OMIM entries to Medical Subject Headings (MeSH) disease categories and assessed enrichment per organ system category. We found that our subnetwork is significantly enriched for genes mutated in mendelian diseases affecting the central nervous system (Fisher’s exact *p = 0.0017*, Table S5), validating our observation of enrichment in fetal brain. We note that this enrichment is not in the inflammatory/immune neurological disease sub-category, suggesting no overlap with the discrete immune signature we found. Samocha *et al* have previously reported that constrained genes are also enriched for *de novo* mutations associated with autism spectrum disorders, further strengthening our conclusion that this constrained subnetwork represents a brain-related biological process.

### The constrained module is preferentially expressed in early brain development

To further elucidate the relevance of our constrained module to brain physiology, we interrogated expression data for multiple brain structures across developmental stages from the BrainSpan project^9^. We found a strong signature of preferential expression in very early stages of development, which declines rapidly and is absent by mid-gestation and remains inactive after birth into adulthood (Figure 3A and Table 3). Several transitional structures in the early brain exhibit significant preferential expression levels, including the ganglionic eminences that eventually form the ventral forebrain and the early structures of the hippocampus. The latter structure shows the most consistent signature across developmental time, with the module’s pattern of expression gradually weakening and becoming non-significant by mid gestation (post-conception weeks 16-18; Figure 3B). These results, taken with the likely involvement of constrained genes in fundamental processes of mitosis and transcriptional regulation, suggest this gene module is relevant to developmental patterning at crucial time points in early brain development.

**Table 3:** the 72-member constrained gene subnetwork is preferentially expressed in a range of tissues and brain structures. We find strong enrichment in a variety of tissues, predominantly neural and immune-derived samples sourced from the Roadmap Epigenome Project (REP) and the BrainSpan Atlas. We report only tissues passing significance with two conservative independent empirical approaches: random permutation of preferential expression values for the subnetwork across tissues (permutation); and comparison to the largest subnetworks detected when we permute constraint scores for all 9729 InWeb genes.

| Source | Tissue | Developmental stage | Permutation<br>p-value | Resampled<br>p-value | Tissue-specific<br>genes |
| --- | --- | --- | --- | --- | --- |
| REP | CD34 | Perinatal (cord blood) | 0.00100 | 0.00100 | 10 |
| REP | Fetal brain | Fetal | 0.01100 | 0.00100 | 16 |
| REP | CD8 | Adult (>20 years) | 0.01700 | 0.00100 | 10 |
| REP | Fetal thymus | Fetal | 0.04800 | 0.00100 | 5 |
| BrainSpan | Caudal ganglionic eminence | 2A (8-9 pcw) | 0.00125 | 0.00125 | 20 |
| BrainSpan | Dorsolateral prefrontal cortex | 2A (8-9 pcw) | 0.00125 | 0.00125 | 16 |
| BrainSpan | Hippocampal anlage | 2A (8-9 pcw) | 0.00125 | 0.00125 | 17 |
| BrainSpan | Lateral ganglionic eminence | 2A (8-9 pcw) | 0.00125 | 0.00125 | 19 |
| BrainSpan | Primary motor-sensory cortex | 2A (8-9 pcw) | 0.00125 | 0.00125 | 20 |
| BrainSpan | Medial frontal cortex | 2A (8-9 pcw) | 0.00125 | 0.00125 | 19 |
| BrainSpan | Orbital frontal cortex | 2A (8-9 pcw) | 0.00250 | 0.00125 | 14 |
| BrainSpan | Parietal neocortex | 2A (8-9 pcw) | 0.00250 | 0.00125 | 18 |
| BrainSpan | Medial ganglionic eminence | 2A (8-9 pcw) | 0.00375 | 0.00125 | 18 |
| BrainSpan | Occipital neocortex | 2A (8-9 pcw) | 0.00500 | 0.00125 | 18 |
| BrainSpan | Hippocampus | 2B (10-12 pcw) | 0.00625 | 0.00125 | 18 |
| BrainSpan | Hippocampus | 3A (13-15 pcw) | 0.00625 | 0.00125 | 19 |
| BrainSpan | Primary somatosensory cortex | 3A (13-15 pcw) | 0.01250 | 0.00125 | 20 |
| BrainSpan | Primary visual cortex | 4 (19-24 pcw) | 0.01750 | 0.00125 | 22 |
| BrainSpan | Posterior superior temporal cortex | 3B (16-18 pcw) | 0.01875 | 0.00125 | 22 |
| BrainSpan | Posteroventral parietal cortex | 3A (13-15 pcw) | 0.02250 | 0.00125 | 19 |
| BrainSpan | Cerebellar cortex | 4 (19-24 pcw) | 0.02500 | 0.00125 | 19 |
| BrainSpan | Primary motor cortex | 3A (13-15 pcw) | 0.02750 | 0.00125 | 19 |
| BrainSpan | Striatum | 3A (13-15 pcw) | 0.04125 | 0.00125 | 17 |
| BrainSpan | Dorsolateral prefrontal cortex | 4A (19-24 pcw) | 0.04625 | 0.00250 | 21 |

**Figure 3:**
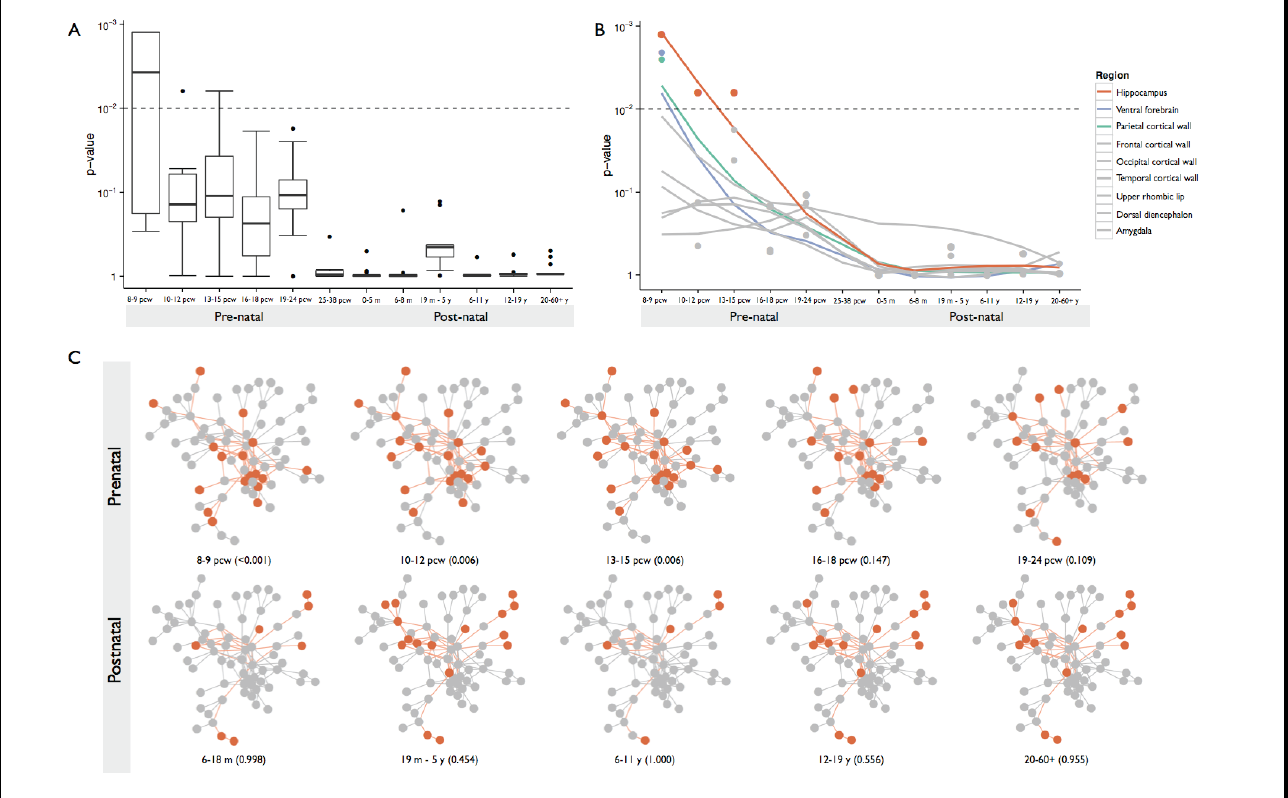
the 72-member selectively constrained gene subnetwork is active in early brain development, particularly in the hippocampus. A: the constrained subnetwork shows elevated signatures of preferential expression in early stages of brain development. B: the signature is most robust in the hippocampus and its ancestral structures (orange), with some enrichment in ventral forebrain and parietal cortical wall structures very early in development (8-9 post-conception weeks). C: The constrained subnetwork shows significant preferential expression in early developmental stages, with patterns of expression losing this enrichment signature by mid-gestation. Overall, these data suggest the constrained subnetwork is specifically active in very early stages of hippocampal formation.

## Discussion

We have shown that selective constraint influences sets of interacting genes involved in core cellular control processes, and that these have elevated expression levels in early stages of central nervous system development. We found the strongest enrichment in the early hippocampal stages at post-conception weeks 8-9, with additional signals in ventral forebrain structures and the parietal cortical wall. This stage of development involves neuronal proliferation through carefully orchestrated sequences of cell differentiation during developmental patterning across the brain. As the constrained subnetwork we have detected is enriched for genes involved in the control of mitosis and transcription, we speculate that it plays a fundamental role in these processes. Our finding that neurological mendelian disease genes are over-represented, combined with previous reports of *de novo* mutations affecting autism spectrum disorders, intellectual disability and epileptic encephalopathy, further support this notion, indicating that any perturbation leads to severe phenotype. This strong limitation in tolerance may also explain our observation of enrichment in immune cell populations, as precise control of developmental decisions is crucial to the correct differentiation of the lymphoid and myeloid lineages throughout life. As the selective constraint scores are by design corrected for both coding sequence length and GC bias^4^, constraint is more likely to be due to intolerance of changes to protein function rather than structural characteristics of the encoded proteins.

More broadly, our results present a glimpse into how natural selection may coordinately shape groups of genes. Most studies of selection aim to identify specific alleles inconsistent with the nearly neutral model of drift, with particular success in studies of recent positive selection^12, 13^. We suggest that the majority of these effects represent near-mendelian effects on relevant phenotypes, which are the actual targets of selective forces: for example, variability in lactase persistence is almost entirely explained by any one of handful of necessary and sufficient alleles^14^. However, the majority of human traits are polygenic, and selection would exert far weaker effects on relevant alleles, which only explain a fraction of phenotypic variance. Although such polygenic adaptation^15^ has proven difficult to detect thus far, our data provide confirmation that selective forces can act on groups of genes involved in the same process, supporting the notion that adaptation can act coordinately on multiple genes.

We have presented a robust approach to identifying sets of interacting genes under selective constraint and placing these into biological context, using the wealth of from genome-scale data produced by large-scale public projects. Our approach builds on robust statistical frameworks to interrogate single variants or genes and thus provides previously lacking biological context from which further hypotheses can be drawn. The approach is flexible and not restricted to studies of constraint: measures of other forms of natural selection, non-human hominid introgression, common and rare variant disease association and any other gene-wise measures can be analyzed in our framework. Further, as PINTS is modular, appropriate tissue atlases can be used to meaningfully interpret results. We believe our work represents a new class of approaches that can leverage multiple genome-scale datasets to gain new insight into biological activities responsible for health and disease.

## Materials and methods

### Selective constraint data

We have used selective constraint scores as previously described^4^. Briefly, we used a mutation rate table—containing the probability of every trinucleotide XY_1_Z mutating to every other possible trinucleotide XY_2_Z—based on intergenic SNPs from the 1000 Genomes project and the sequence of a gene to determine that gene’s probability of mutation. These sequence context-based probabilities of mutation were additionally corrected for regional divergence between humans and macaques as well as the depth of coverage for each base in an exome sequencing study. Given the high correlation (Pearson’s r = 0.94) between the probability of a synonymous mutation in a gene with the number of rare (MAF < 0.01%) synonymous variants in that gene seen in the NHLBI’s Exome Sequencing Project, we used a linear model to predict the number of rare missense variants expected per gene in the same dataset. The difference between observation and expectation was quantified as a signed Z score of the chi-squared deviation. The missense Z score was used as the basis for determining selective constraint. In this study, we took a conservative approach to assessing selective constraint, using the Bonferroni correction for number of InWeb genes to derive a significance threshold of *p*_*c*_ < *5 × 10*^*−6*^.

### Detecting selectively constrained subnetworks in protein-protein interaction data

We used InWeb, a previously described comprehensive map of protein-protein interactions, containing 169,736 high-confidence interactions between 12,687 gene products, compiled from a variety of sources^10^. By mapping ENSEMBL IDs, we were able to identify 9729 interconnected genes with constraint scores from Samocha *et al* also present in the REP expression data (below), to which we restricted our analysis.

To detect clusters of interacting constrained genes, we used a heuristic form of the prize-collecting Steiner tree (PCST) algorithm^16, 17^, which has been previously applied to protein-protein interaction data^11^. The canonical form of the PCST algorithm takes a connected, undirected graph *G(V,E,w,u)* with *V* vertices and *E* edges, with vertex weights *w* and edge weights *u*; it then finds the connected subgraph *T(V’,E’)* with maximal *profit(T)*, which is some function of *(w’-u’)*. By definition, *T* is a minimal spanning tree. The algorithm thus identifies the set of nodes with the strongest signal given the *cost* of their connecting edges. The classical PCST algorithm is, however, *NP-hard*, which makes it computationally intractable on the scale of InWeb^16^. Several heuristic simplifications have been proposed, including one previously validated as suitable for protein-protein interaction networks^11^. This approach partitions the set *V* into *null* (with weights *w < 0*) and *signal* (with weights *w > 0*) vertices (genes) and equal edge weights *e* before searching for *T*. Beisser *et al* have implemented this approach in the BioNet package for the R statistical language^18^. Here, we define signal genes as those with constraint scores passing the Bonferroni threshold of *p*_*c*_ < *5 × 10*^*−6*^, and calculate the weights as *w* = − *log*(*p*_*c*_) + *log*(*5 × 10*^*−6*^). The PCST algorithm returns a single, maximal *T* solution; to discover further independent subnetworks, we apply the method iteratively after we assigning gene nodes in the previously discovered solution to be null.

The algorithm always returns a solution for *T*, so we sought to assess the significance of our observations empirically. To understand if the observed solution is unlikely by chance, we permuted the constraint scores of genes 1000 times and for each iteration ran the heuristic PCST to generate 1000 random *resampled subnetworks* (these are also used in the tissue-specificity analyses described below). We then quantified the following key parameters and assessed how many random subnetworks had values exceeding those of the true discovered subnetwork: size (number of gene nodes); density (number of connections); clustering coefficient and total amount of constraint explained (sum of constraint scores).

### Gene expression data processing and preferential expression analysis

We obtained gene expression data for a cosmopolitan set of tissues from the Roadmap Epigenome Project (REP)^8^. The REP data consists of 88 samples across 27 tissue types from diverse human organs, profiled on the Affymetrix HuEx-1_0-st-v2 exon array, which we downloaded on 9/25/2013 from http://www.genboree.org/EdaccData/Current- Release/experiment-sample/Expression_Array/. We processed these data using standard methods available from the BioConductor project^19, 20^. Briefly, we removed cross-hybridizing probesets, applied RMA background correction and quantile normalization and then summarized probesets to transcript-level intensities. We then mapped transcripts to genes using the current Gencode annotations for human genes (version 12). Transcripts with no match in Gencode were removed and the remaining transcripts we again quantile normalized. We then assigned transcript expression levels to their matching genes. Where multiple transcripts mapped to the same gene we used the transcript with maximum expression over all cell types.

The Brainspan atlas^9^ data are available as processed, gene-level expression levels from from http://www.brainspan.org/static/download.html. We mapped these genes to the InWeb gene set using ENSEMBL IDs, and quantile normalized data for the overlapping genes. We then grouped replicate data by developmental stage and brain structure and calculated preferential expression as described above.

We used a previously described approach to detect tissue-specific expression across each tissue atlas^21^. Briefly, we group together replicates from the same cell type and compute pairwise differential expression between all pairwise combinations of tissues, using an empirical Bayes approach to account for variance shrinkage^22^. Thus, for each gene there are 26 linear model coefficients and associated *p* values for each tissue, quantifying the comparison to all other tissues. For each gene in each tissue, we then capture the overall difference in expression from all other tissues as the sum of these coefficients. To reduce noise, only coefficients with *p < 0.0019* (*p < 0.05* with Bonferroni correction for 26 tissues) are considered. Rescaling all coefficient sums across all genes values to the range [-1,1] gives us a final preferential expression score. Intuitively, a gene highly expressed in only one tissue would get a high positive enrichment score in that tissue, as it is differentially expressed compared to all other tissues. The score is directional, strong negative values indicate very low expression in one tissue compared to all others. We partition the overall distribution into deciles and define preferential expression in a tissue if a gene has a score > 0.1.

### Scoring subnetwork tissue specificity

To score the tissue specific expression of a subnetwork, we detect which genes in the subnetwork are preferentially expressed in each tissue of our expression atlas and assess the joint probability of this observation. To do so correctly we must account for the connections between genes and the pattern of preferential expression of each gene across the tissue atlas. Formally, we consider the subnetwork as a Markov random field with a particular configuration of preferentially expressed nodes in each atlas tissue. We compute a score for each configuration using a standard scoring function^23^:

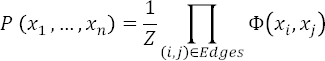

The partition function Z is defined as:

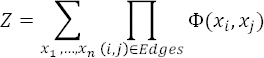

where *x*_*i*_ (*i* = 1,…, *n*) represents a binary tissue specificity of the genes in the subnetwork for a given tissue with values either 1 (expressed) or 0 (not expressed). The Φ(*x*_*i*_, *x*_*j*_) factor lists the co-occurrence of two connected nodes across tissues. This is calculated from the thresholded preferential expression data, and each pair of connected nodes is assigned exactly one *configuration* in each tissue, so that

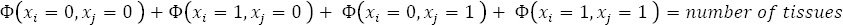

We assess the significance of these scores using two conservative permutation approaches. First we assess how likely we are to see each observed configuration (i.e. each pattern of detected/not detected nodes) in each tissue of the atlas. We do this by permuting the preferential expression scores across tissues for each gene independently and rescoring the configuration found in each tissue. This alters the co-expression structure across genes and empirically assesses how likely we are to see a particular configuration of a specific subnetwork by chance. Second, we estimate the probability of observing the extent of tissue specificity in each tissue. We construct the null expectation by scoring the *resampled subnetworks* generated by permutation above in each tissue and compute the empirical significance from this distribution of scores.

To ensure our results are not artifacts of a specific preferential expression threshold, we repeat this analysis across a spectrum of preferential expression thresholds (See Table S3).

### Pathway analysis

To test if any biological pathways are over represented in a subnetwork, we use the Gene Set Enrichment Analysis (GSEA) approach ^24^. We obtained the full list of curated canonical pathways from the GSEA website (http://www.broadinstitute.org/gsea/msigdb/collections.jsp) and mapped the 9729 genes to each pathway using HUGO IDs. We then test for enrichment of subnetwork members over background using the hypergeometric test.

### Online Mendelian Inheritance in Man (OMIM) analysis

To test if genes in the subnetwork are more likely to harbor pathogenic mutations causing Mendelian diseases than expected by chance, we retrieved OMIM records for all 9729 genes using the biomaRt package in BioConductor^20^. We then tested whether the proportion of 107 subnetwork genes with OMIM entries was higher than the background proportion of the full set of 9729 in our analysis using Fisher’s exact test (Table S4). We then mapped all OMIM entries to Medical Subject Headings (MeSH) disease categories using the Comparative Toxicogenomics Database (CTD) MEDIC disease vocabulary^25^ and assessed enrichment in any disease category, again using Fisher’s exact test (Table S6).

## Acknowledgements

We acknowledge our use of the gene set enrichment analysis, GSEA software, and Molecular Signature Database (MSigDB), available at http://www.broad.mit.edu/gsea/

